# Analysis of *Drosophila* and mouse mutants reveals that Peroxidasin is required for tissue mechanics and full viability

**DOI:** 10.1101/2023.07.19.549730

**Authors:** K. Elkie Peebles, Kimberly S. LaFever, Patrick S. Page-McCaw, Selene Colon, Dan Wang, Aubrie M. Stricker, Nicholas Ferrell, Gautam Bhave, Andrea Page-McCaw

## Abstract

Basement membranes are thin strong sheets of extracellular matrix. They provide mechanical and biochemical support to epithelia, muscles, nerves, and blood vessels, among other tissues. The mechanical properties of basement membranes are conferred in part by Collagen IV (Col4), an abundant protein of basement membrane that forms an extensive two-dimensional network through head-to-head and tail-to-tail interactions. After the Col4 network is assembled into a basement membrane, it is crosslinked by the matrix-resident enzyme Peroxidasin to form a large covalent polymer. Peroxidasin and Col4 crosslinking are highly conserved, indicating they are essential, but homozygous mutant mice have mild phenotypes. To explore the role of Peroxidasin, we analyzed mutants in Drosophila, including a newly generated catalytic null, and found that homozygotes were mostly lethal with 13% viable escapers. A Mendelian analysis of mouse mutants shows a similar pattern, with homozygotes displaying ∼50% lethality and ∼50% escapers. Despite the strong mutations, the homozygous escapers had low but detectable levels of Col4 crosslinking, indicating that inefficient alternative mechanisms exist and that are probably responsible for the viable escapers. Further, fly mutants have phenotypes consistent with a decrease in stiffness. Interestingly, we found that even after adult basement membranes are assembled and crosslinked, Peroxidasin is still required to maintain stiffness. These results suggest that Peroxidasin crosslinking may be more important than previously appreciated.

## Introduction

The basement membrane is a distinct sheet-like type of extracellular matrix found in virtually all animals, in nearly all tissues. Basement membranes underlie epithelia, and they surround muscles, nerves, blood vessels, and organs. In terms of function, basement membrane acts as a mechanical support, a reservoir of signaling molecules, and a signaling insulator (Jayadev and Sherwood, 2017; Ramos-Lewis and Page-McCaw, 2018). Basement membranes are comprised of independent networks of laminin and type IV collagen, with other conserved proteins and proteoglycans assembling on these networks (Pastor-Pareja and Xu, 2011; Ramos-Lewis et al., 2018). The most abundant proteins in basement membranes are type IV collagen heterotrimers, and these form a branched network by assembling with each other at their N- and C-termini, with four heterotrimers assembling at the N-terminal 7S domain and two heterotrimers assembling at the C-terminal NC1 domain. The collagen network determines much of the mechanical strength of the basement membrane because after collagen is assembled in basement membranes, it is crosslinked to form a continuous large covalent network.

The NC1 domain is crosslinked through a covalent bond that appears unique for this purpose, a sulfilimine bond (S=N) that joins a hydroxylysine of one NC1 domain to a methionine domain of another (Vanacore et al., 2009). Sulfilimine crosslinking is accomplished by the enzyme Peroxidasin, first discovered in Drosophila as an enzyme that localized to the basement membrane (Nelson et al., 1994), a location that is consistent with its function in crosslinking assembled basement membrane. Peroxidasin is not known to have functional substrates other than the NC1 domain (He et al., 2020), and like collagen IV, it is highly conserved throughout the animal kingdom (Fidler et al., 2014). Together, these observations suggest that it would be an essential enzyme.

Given this expectation, it was surprising then when the first mouse mutant in *Peroxidasin*, *Pxdn^T3816A^*, was reported to be homozygous viable with eye defects, anterior segment dysgenesis and micropthalmia, as the only prominent phenotype (Yan et al., 2014). A second mutation was generated by CRISPR, deleting exon 1, and this mutant also had small eyes (Kim et al., 2019). Indeed, human alleles of *Pxdn* have been identified in families with inherited eye defects (Choi et al., 2015; Khan et al., 2011). These consistent phenotypes, apparently restricted to the anterior chamber of the eye, suggested that Pxdn had a limited role in matrix stability. A third mouse mutation in *Pxdn* was generated by homologous recombination, and homozygotes were tested directly for basement membrane stiffness. In line with its molecular function, Pxdn^K01^ basement membranes were significantly and substantially less stiff in a tensional stiffness assay (Bhave et al., 2017).

*Peroxidasin* mutants also exist in the fruitfly *Drosophila melanogaster* (where it is abbreviated *Pxn*) although there is no comprehensive report on *Pxn* mutant phenotypes. Because of the high conservation in sequence between fly and mouse Peroxidasin, the *Drosophila* mutants may be useful for understanding its function. We begin this study by analyzing the existing fly mutants, generate a new catalytic null mutation in fly, and compare its phenotype with two mouse knockout alleles. We find that in both species, *Peroxidasin* homozygotes are usually lethal, but even for the strongest catalytic null allele, some homozygous progeny survive. The fly phenotypes confirm the role of Pxn in tissue stiffness, as homozygotes die during life stages with large mechanical strain, hatching and pupal eclosion. Further, adult flies have dysmorphic gut muscles. *Peroxidasin* mutants also have more fragile basement membranes, demonstrated in an osmotic swelling assay. Finally, we use a mechanical stiffness assay to demonstrate that even after basement membranes are assembled and cross-linked in wild-type adult animals, continuous Pxn function is required to maintain tissue stiffness.

## Results and Discussion

### Fly *Peroxidasin* mutants are lethal with incomplete penetrance

The Peroxidasin protein is ancient and highly conserved (Fidler et al., 2014), so it is surprising that *Peroxidasin* (*Pxdn*) mutant mice are viable, with the only obvious phenotype being severe eye malformations (Kim et al., 2019; Yan et al., 2014). To better understand the functions of Peroxidasin, we analyzed the lethality of two existing *Drosophila Peroxidasin* (*Pxn*) mutants, *Pxn^MI01492^* and *Pxn^f07229^.* Both alleles are recessive loss-of-function, and they are caused by the insertion of two different transposons in two different *Pxn* introns (Fig. 1A). In these mutants as in other transposon alleles, the coding region remains intact, and the mutation is caused by a reduction in gene product levels. Analyzing the progeny of heterozygous crosses, we counted a very small number of viable homozygotes for each allele, and we inferred that 3.6% (*MI01492*) or 2.5% (*f072229*) of *Pxn* homozygous mutants were viable (Fig. 1B,C). However, when these insertion alleles were placed *in trans* so that a fly had one of each allele, viability improved: 13% of *Pxn^MI01492/f07229^* mutants were viable (Fig. 1D). Because background effects were reduced in the trans-heterozygotes, 13% viability is a better reflection of the lethal phenotype of these *Pxn* mutants.

**Fig. 1:**
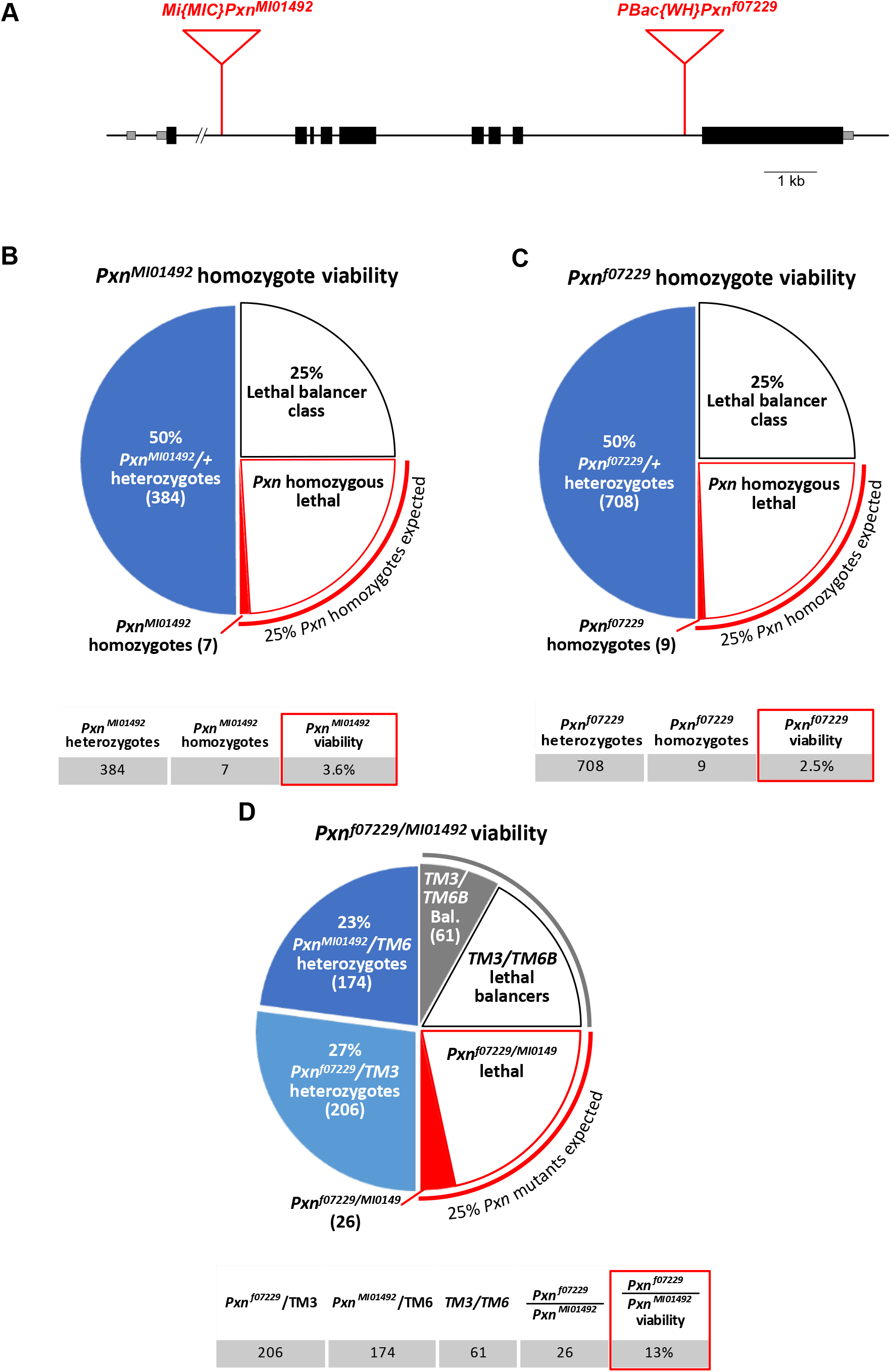
Fly *Pxn* inserHonal mutants are parHally viable.

We hypothesized that the 13% viability of the trans-heterozygotes reflected residual function of the insertional alleles, and we expected that a *Pxn* catalytic null mutant would be a fully penetrant lethal mutation. Further, we expected *Pxn* null mutants to be embryonic lethal because basement membrane is first assembled in embryos (Lunstrum et al., 1988; Matsubayashi et al., 2017). To generate a null mutation, we targeted CRISPR guide RNAs to the sequence encoding the catalytic domain. The resulting mutation, *Pxn^11^*, is a 727 bp deletion that generates a frame-shift, removing the catalytic core of the Peroxidasin enzyme starting at N1041 (Fig. 2A,B). Contrary to our expectations, the *Pxn^11^* catalytic null mutant has similar viability as the *Pxn^MI01492/f07229^* transheterozygotes: 13% of the homozygous animals survived while 87% of them were lethal (Fig. 2C).

**Fig. 2:**
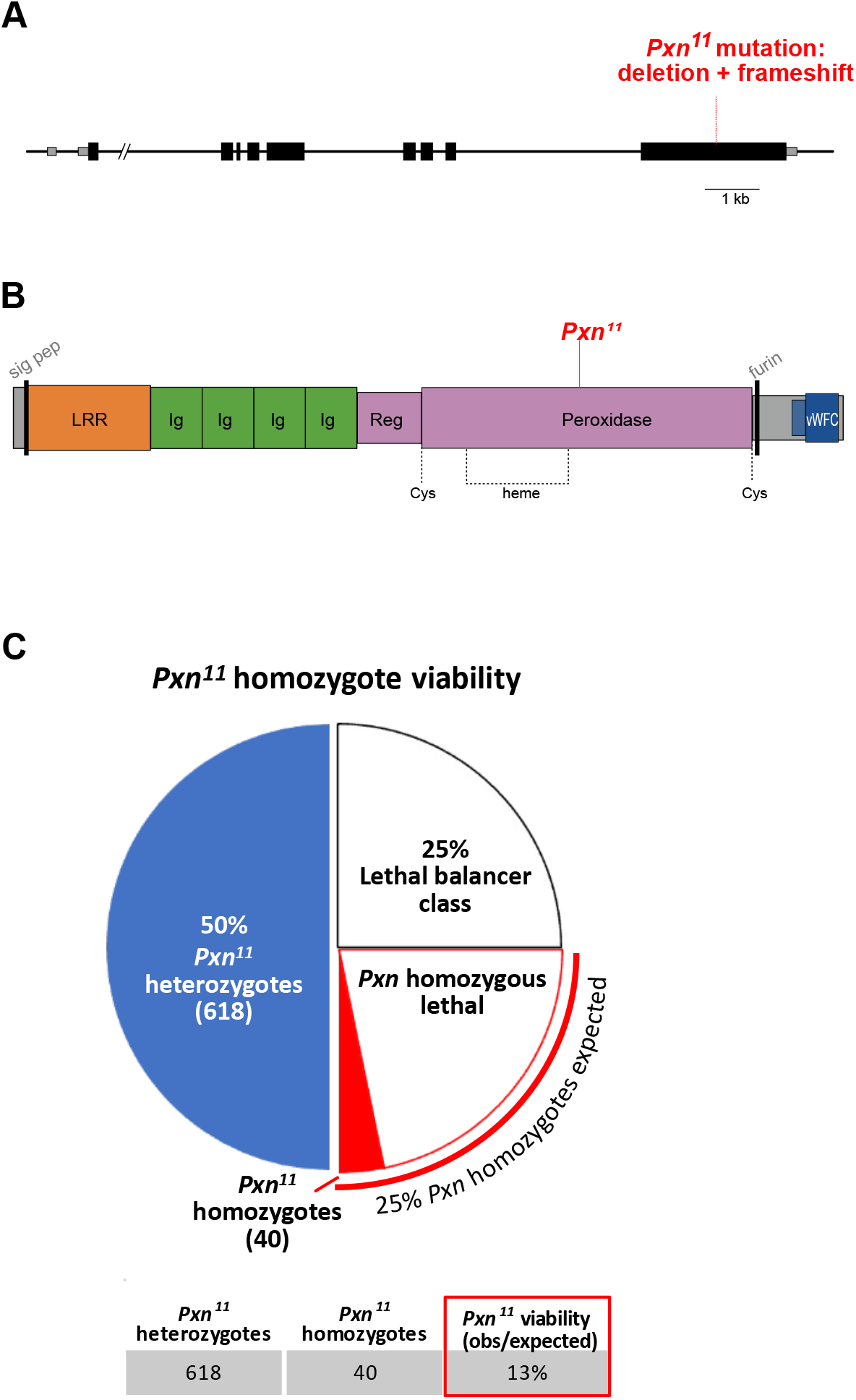
Fly *Pxn¹¹* mutant, a catalyFc null, is parFally viable.

### Mouse *Peroxidasin* mutations are lethal with incomplete penetrance

Given the fly catalytic null mutants were mostly lethal with some adult survivors, we wondered if there could be a similar range of viable/lethal progeny in mice. To address this question, we generated two mutant alleles at the mouse *Pxdn* locus. The first allele, designated *Pxdn^K0^*^1^, was generated by homologous recombination of the KOMP vector into the *Pxdn* gene locus, replacing the 9^th^ exon with sequences that generate a gene-trap by splicing to LacZ followed by polyadenylation sequences before the exon 9 sequence (Skarnes et al., 2011). *K01* generates a predicted Pxdn protein that is truncated in the first Ig domain of Pxdn, which would have no catalytic function (Fig. 3A,B). *K01* was generated in B6/129 hybrid embryonic stem cells, and it was outcrossed separately into C57BL/6 and 129S2 backgrounds for 10 generations. In both backgrounds, viable homozygous mutant *Pxdn^K01/K01^* adults were observed, and they were blind with a defect in development of the anterior chamber of the eye, as previously described for this and other *Pxdn* alleles (Bhave et al., 2017; Kim et al., 2019; Yan et al., 2014). To investigate whether *Pxdn* mutants were fully viable, we genotyped all the pups from multiple heterozygous crosses to identify the numbers of each progeny class and compared them to Mendelian expectations. Fewer *Pxdn^KO1/KO1^* were detected post-weaning than expected, indicating that 69% of *Pxdn^KO1/KO1^* mutants in the 129S2 background were viable (Figure 3C). A chi-square test found the fraction of observed *Pxdn^KO1/KO1^* pups to be significantly different than the expected Mendelian ratio of 25%, with X^2^= 20.6 with a corresponding p value < 0.0001 (Fig. S1). To determine whether this loss in viability was strain specific (i.e. caused by interactions with modifier alleles of other genes), the mutants were outcrossed to the C57BL/6 background and viability was again measured: in the C57BL/6 background only 46% of *Pxdn^KO1/KO1^* mutants were viable.

**Fig. 3:**
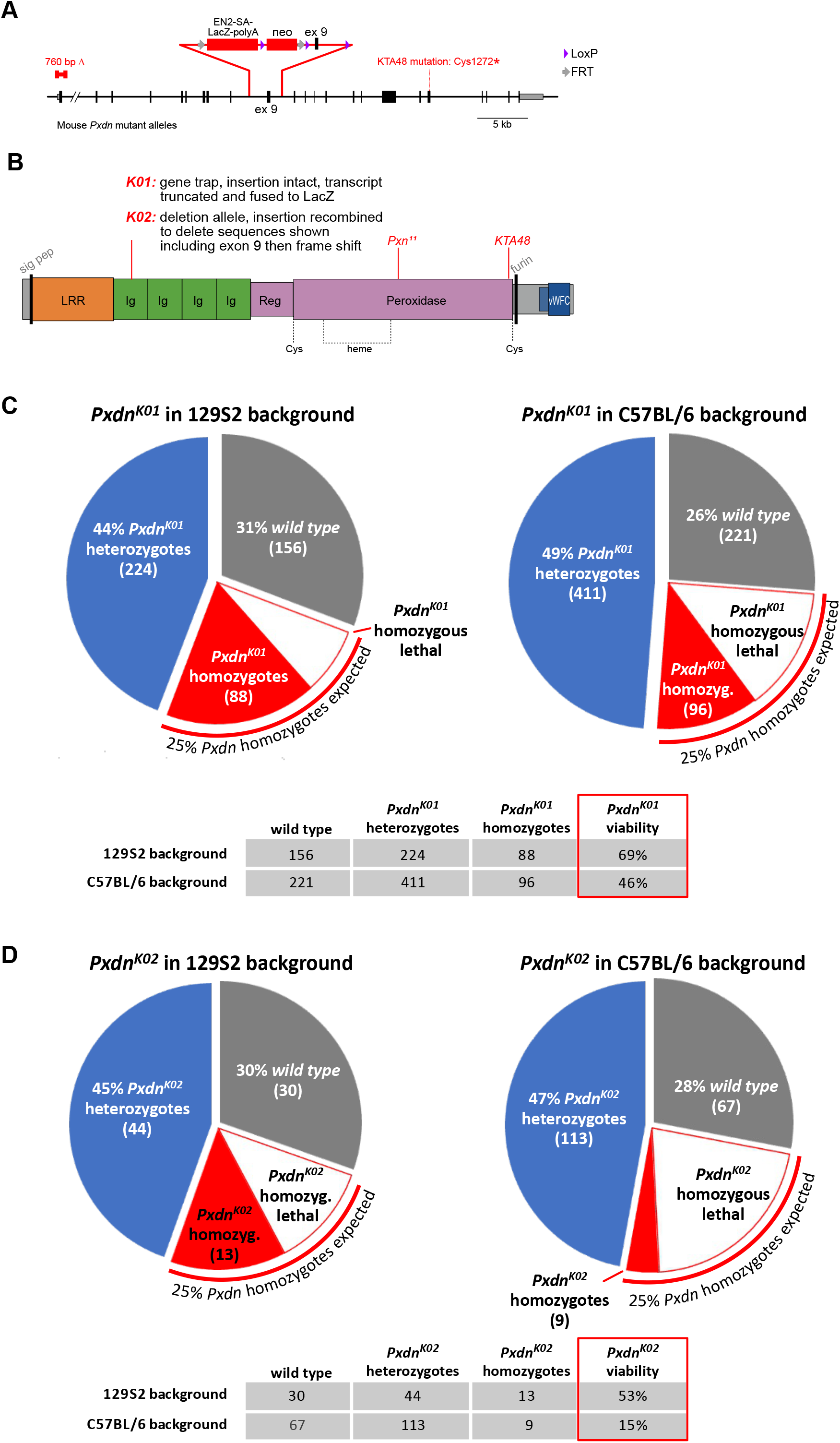
Mouse Pxdn homozygotes are parMally viable.

To determine whether the partial viability may be due to leakiness of the *KO1* allele, we generated a second allele designated *Pxdn^KO2^*, excising most of the insertion sequence including exon 9 sequence by crossing sequentially to Flp and Cre recombinase lines. The *Pxdn^K02^* allele was outcrossed separately into C57BL/6 and 129S2 backgrounds for 10 generations. The *KO2* homozygotes were 53% viable in the 129S2 background; in the C57BL/6 background, *K02* had further reduced viability, with only 15% of expected homozygotes surviving through weaning (Fig. 3D). For all four combinations of alleles and backgrounds, viability was significantly reduced from the 25% expectation as determined by chi-square test (Fig. S1). We conclude that, for both flies and mice, strong mutations in the *Peroxidasin* gene locus significantly reduce viability. In principle, the partial penetrance of the lethal phenotype could be caused by a leaky mutation with residual function, or it could reflect a second source of variable low-level crosslinking. Because the fly mutation deletes the catalytic core of the protein, we ruled out the possibility of leaky function and investigated the levels of crosslinking in these alleles.

### Fly and mouse *Peroxidasin* mutants retain low levels of collagen IV NC1 crosslinking

Peroxidasin generates sulfilimine crosslinks between two head-to-head collagen IV NC1 domains in the basement membrane. The enzyme catalyzes the formation of hypobromous acid (HOBr) from bromide and hydrogen peroxide, and HOBr reacts with the uncrosslinked sites in an NC1 hexamer to generate sulfilimine bonds between Met93 of one NC1 and Hyl211 of the other, creating an NC1 dimer (McCall et al., 2014). Because each NC1 domain contains both Met93 and Hyl211, a dimer can have one or two sulfilimine crosslinks, called D1 and D2, which migrate with different mobilities on polyacrylamide gel electrophoresis (McCall et al., 2014). To investigate the extent of NC1 crosslinking in the new fly and mouse *Peroxidasin* mutants, we analyzed the level of D1, D2 and uncrosslinked monomers. We treated lysates from *Pxn^11^* whole adult flies with collagenase to release the NC1 domain from the insoluble basement membrane and then probed a western blot with a low-affinity antibody raised against fly NC1 (McCall et al., 2014). The D2 double-crosslinked dimer was not observed, but the levels of D1 indicated there was about 11% of the total crosslinking in *Pxn^11^* mutant adults compared to wild-type (Fig. 4A). This is similar to the level of crosslinking previously observed in mouse *K01* mutants, which have about a third of the level of crosslinking with virtually no D2 dimers (Bhave et al., 2017). For the new *K02* allele, D2 dimers were not observed, but some D1 crosslinking was observed (Fig. 4B). These results suggest that NC1 D1 crosslinking occurs at low levels even without Peroxidasin, perhaps reacting with the hypohalous acid products of myeloperoxidase or other related heme peroxidases, which exist in both mouse and fly. Alternatively, it is possible there is a low level of uncatalyzed reaction; for most enzymes this on-rate is too low to be detected given turnover, but in this case the crosslinked collagen products are long-lived, possibly enabling detection. Whatever the source, this low level of crosslinking from an alternative source probably supports the partial viability of *Peroxidasin* mutants observed in both species. Thus, it may be reasonable to think of Peroxidasin as a specialist enzyme that promotes consistent and efficient crosslinking rather than the variable low-level of crosslinking observed in its absence.

**Figure 4:**
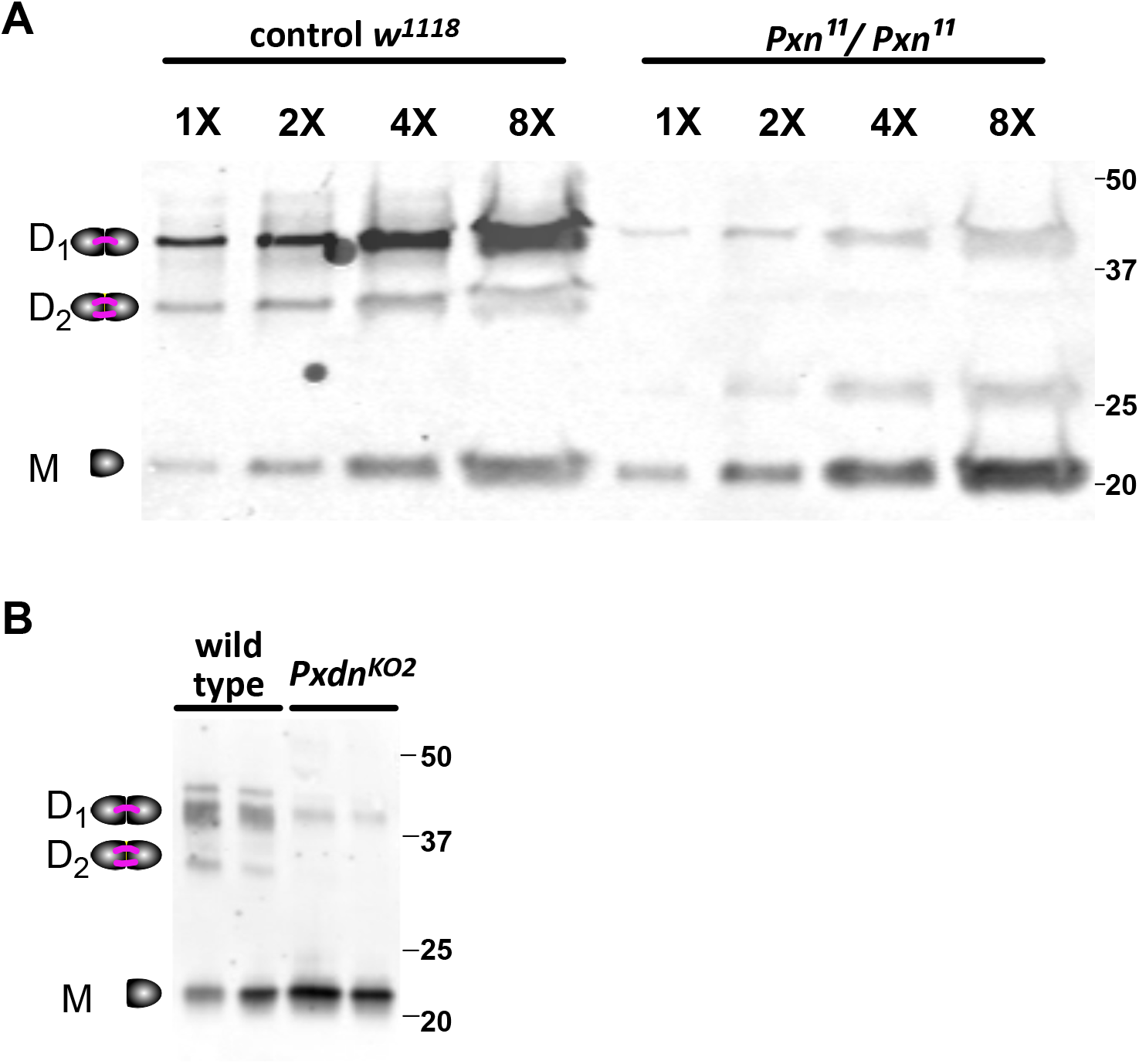
Collagen IV NC1 crosslinking is reduced but not absent in fly and mouse mutants of *Peroxidasin*. **A.** Western blot of whole adult flies homozygous for the *Pxn11* catalytic null mutant shows 11% of the wild-type levels of NC1 domain crosslinking. Crosslinking was probed by anti-NC1 staining of the insoluble fraction of fly lysate. **B.** Western blot of whole mouse kidneys shows reduced NC1 crosslinking in *Pxdn^K02^* homozygotes compared to wild-type. Crosslinking was probed with H22 antibody gainst the NC1 domain of Col4a2.

### *Peroxidasin* mutants die at developmental periods when tissues exert force

To gain a better understanding of the requirements for Peroxidasin, we analyzed *Drosophila Pxn^11^* mutants to determine when they died during development. The *Drosophila* life cycle comprises two distinct body plans, a worm-like larval body plan that develops during embryogenesis, and an adult fly body plan that develops during metamorphosis. Before metamorphosis, the larva grows through three distinct periods or instars, which can be recognized morphologically. We followed the development of hundreds of animals of three genotypes, *Pxn^11^* homozygotes, *Pxn^11^/TM3* (heterozygous control for background effects), and *w^1118^* (control with wild-type *Pxn*), noting their progression to the next stage of development. Although controls had some lethality before hatching, we found that 25-30% fewer *Pxn^11^* mutant embryos hatched compared to controls, a significant effect (Fig. 5A). The larval transitions were similar between *Pxn* mutants and controls (Fig. 5B,C), followed by a slight, but significant, reduction in the ability of mutants to enter metamorphosis (pupariation, Fig. 5D). The largest lethal stage of *Pxn* mutants was observed at the transition from pupa to adult (eclosion), with *Pxn^11^* pupae eclosing at only 25% the rate of controls (Fig. 5E). The developmental lethality data is summarized in a survival curve in Fig. 5F. To investigate pupal lethality further, we observed the morphology of heterozygous and homozygous pupae and found that all *Pxn* homozygous animals, even those that failed to eclose, had fully formed adult bodies visible inside the pupal case (Fig. 5G).

**Figure 5:**
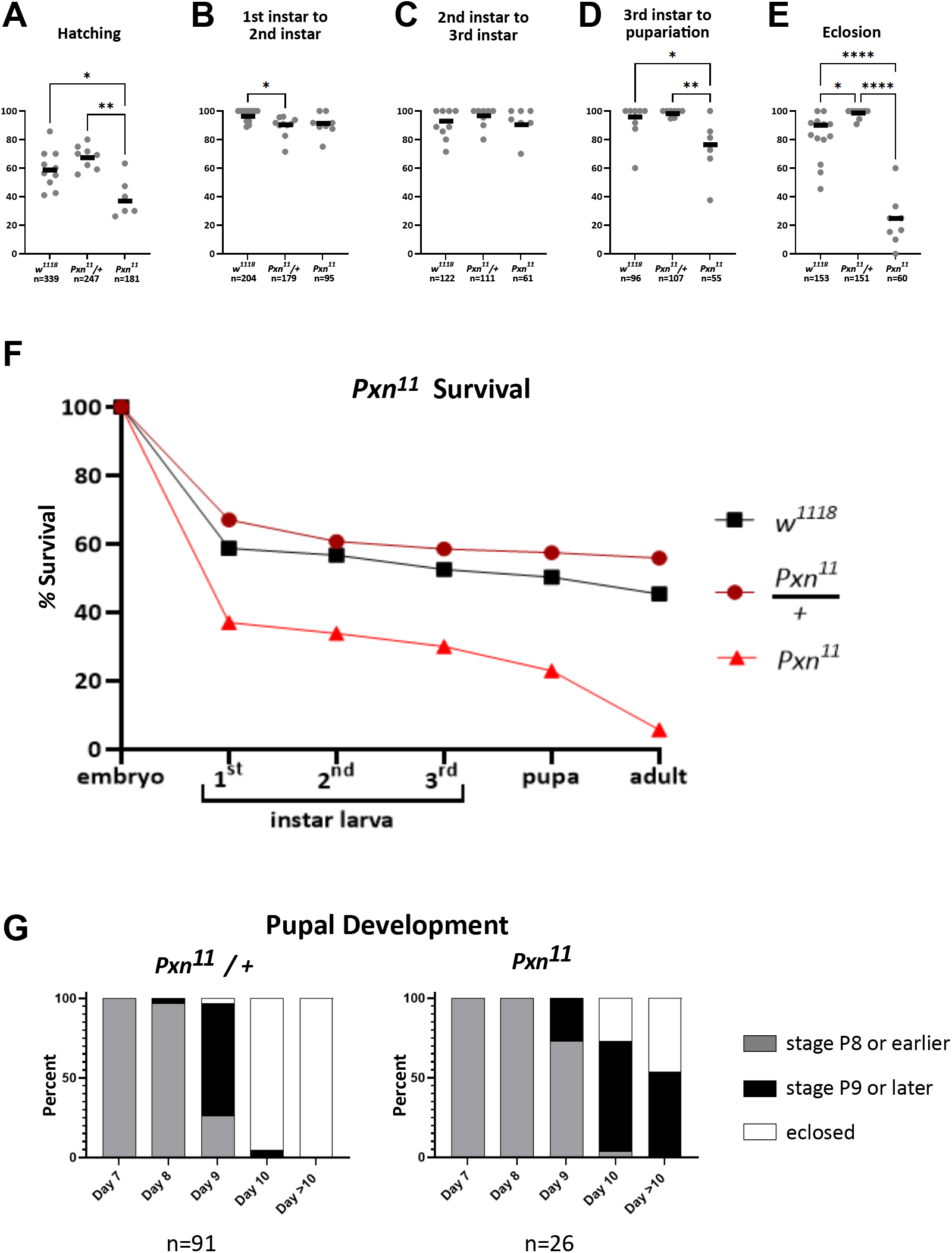
P*x*n11 mutants die at developmental stages that require mechanical force. **(A-E)** Animals of indicated genotype were tracked across each developmental transition. Each individual data point is a % survival within a subpopulation (single vial). The bar represents the cumulative survival of the total number animals. *Pxn11* mutants die most frequently during embryogenesis (A) and eclosion (E), both stages in which the animal must use force to break through either the eggshell or the pupal case. **(F)** Cumulative survival of animals shown in panels A-E. **(G)** *Pxn11* mutant pupae appeared fully developed but could not escape the pupal case. *Pxn11* mutant pupae older than 10 days never eclosed.

Thus, *Pxn* mutants die most frequently during embryogenesis/hatching and eclosion. These stages require mechanical force, as embryos and pupae both push through and break out of the enclosing cuticle, either eggshell or pupal case. Prior to both hatching and eclosion the animal generates large scale highly patterned motor programs which may place the tissues under considerable tension. The conclusion that *Pxn* mutants are less able to exert or withstand physical force is consistent with our understanding of *Pxn* as a Collagen IV crosslinking enzyme.

We tried to analyze fertility in the surviving male and female homozygous Pxn11 mutants, and our initial observations were that they were almost completely sterile. However, we found that the sterility phenotype changed rapidly over the course of three months: testing laying frequency in small groups of homozygous females, we found that flies from later generations laid dramatically more eggs, at about the same rate as controls (Fig. S2). These results indicate that Pxn11 mutants were under selection to pick up suppressors of sterility, and that suppressors were easily generated suggesting multiple genetic targets. The existence of readily available suppressor mutations or conditions may also explain why some Pxn11 mutants survived to adulthood.

Another unexpected genetic interaction was identified when we tried to cross *Pxn^11^* to VkgGFP, a functional Col4a2 molecule tagged at the endogenous locus with GFP before the 7S domain. We were unable to complete this cross because we could not generate female flies heterozygous for *Pxn^11^* and VkgGFP, despite multiple attempts. This genetic interaction suggests that protein interactions at the 7S domain, where Col4 molecules become crosslinked to form dodecamers, may become especially important once crosslinking at the NC1 domain is severely compromised.

### *Peroxidasin* mutants are less stiff and fail under tensile strain

The requirement for *Pxn* during stages that require force suggested that *Pxn* mutants have altered tissue mechanics. Indeed, we previously showed that Pxn^K01^ kidney tubules have reduced stiffness in response to stretching, using an *ex vivo* assay to measure tensile stiffness (Bhave et al., 2017). In flies, we previously observed that the loss of *Pxn* by RNAi-based knockdown caused defects in muscle shape (Howard et al., 2019), and we confirmed that phenotype in *Pxn^11^* homozygous mutants: on dissection of the adult midgut, we observed that the gut peristalsis muscles were dysmorphic (Fig. 6A,B), consistent with a mechanical role of basement membranes in distributing muscle force, an insight from research into muscular dystrophy (Campbell, 1995).

**Fig. 6.**
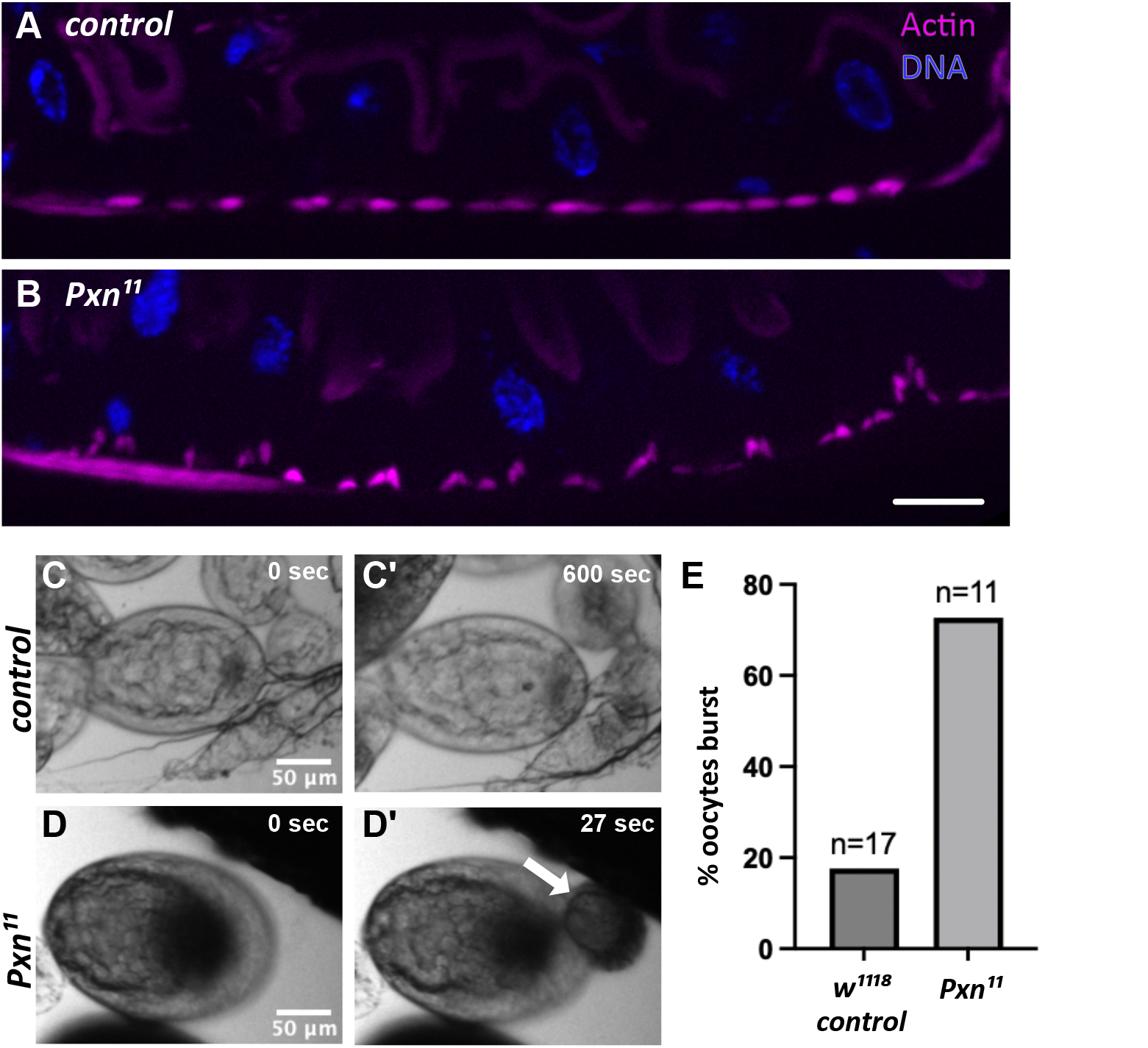
*Pxn^11^* homozygous mutants have defects in tissue mechanics. **(A-B)** Peristalsis muscles of the adult midgut are dysmorphic in the *Pxn*^11^ mutant. Muscles are identified by actin staining in control *w^1118^*(A) and *Pxn*^11^ homozygous mutants (B). Scale bar: 10 µm. **(C-E)** *Pxn*^11^ oocytes burst more frequently (C) than control w1118 oocytes (D) when stretched by osmotic swelling. Oocytes (stages 6-8) were placed in water for 10 minutes and observed for bursting. 18% of control w1118 oocytes burst whereas 73% of *Pxn*^11^ mutants burst (E).

Another basement membrane mechanics assay that has been utilized in *Drosophila* is the oocyte bursting assay. Oocytes develop inside a collagen-IV basement membrane, and when immature oocytes are dissected and submerged in hypo-osmotic water *ex vivo*, the oocytes swell, placing the basement membrane under mechanical strain. After several minutes in hypo-osmotic media, oocytes may burst; the frequency and timing of bursting has been correlated with basement membrane stiffness as measured by atomic force microscopy (Crest et al., 2017). Within 10 minutes in water, *Pxn^11^* mutants burst more frequently than control *w^1118^* oocytes: whereas only 18% of controls burst within 10 minutes, 73% of *Pxn^11^* mutants burst (Fig. 6C-E). This bursting rate is consistent with previous reports of oocyte bursting in basement membrane knockdowns (Crest et al., 2017). Thus, developmental phenotypes, muscle phenotypes, previously reported tensile stiffness, and oocyte bursting rates all indicate that Peroxidasin mutants have altered basement membrane mechanics.

### Continued Peroxidasin-mediated crosslinking maintains the stiffness of adult basement membranes

The fly and mouse mutants clarify that Peroxidasin establishes basement membrane stiffness in development, but they cannot distinguish whether Peroxidasin continues to be required in adults. We previously found that knocking down *Pxn* in adult flies phenocopied a muscle defect related to loss of stiffness, suggesting a continuing requirement for Pxn in maintaining basement membrane stiffness (Howard et al., 2019). These results were surprising, as it is not known that Pxn would be required once the basement membrane was assembled and crosslinked in an adult animal, and Col4 is known to be a long-lived (Decaris et al., 2014; Liu et al., 2019; Liu et al., 2020). To investigate this possibility with a quantitative assay, we fed wild-type flies the Peroxidasin catalytic inhibitor Phloroglucinol (PHG) for 5 days, then assessed the tensional stiffness of their Malpighian (renal) tubules *ex vivo* (Fig. 7). Malpighian tubules, like kidney tubules in mouse, are simple epithelial tubes surrounded by basement membrane.

**Figure 7:**
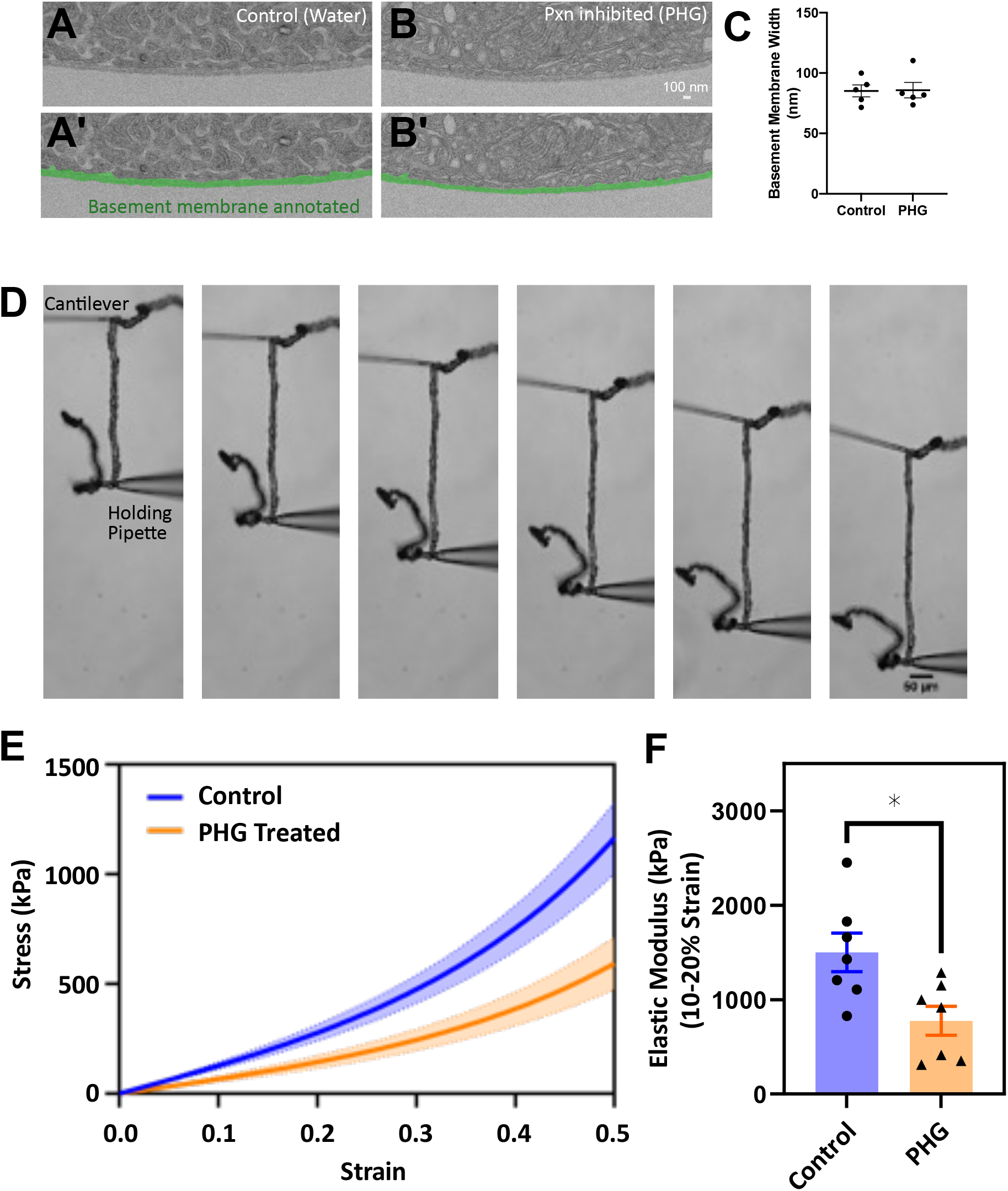
Continued Peroxidasin crosslinking maintains basement membrane stiffness in adult tissues. Malpighian (renal) tubules were dissected from adult wild-type flies that had been fed the Peroxidasin catalytic inhibitor phloroglucinol (PHG) for 5 d prior to dissection. **A-C.** Transmitted electron micrograph of Malpighian tubule basement membrane shows that tubule basement membrane thickness is unchanged after 5 d Peroxidasin inhibition. **D.** Tensile strain assay: Cantilever-based stiffness measurement system. Tubules were stretched between a force calibrated microcantilever and a rigid holding pipette. **E.** Stress-strain curves were calculated from the bending of the cantilever and displacement of the holding pipette. **F.** Elastic moduli of tubules is reduced after 5 d of Peroxidasin inhbition by PHG. Elastic modulus was calculated from slope of the stress-strain curves at 10% and 20% strain. Data were analyzed by unpaired t-test in GraphPad, *p<0.05. See also Fig. S3.

Tubule stiffness is determined by their basement membrane, as stiffness measurements are not significantly different when cells are lysed before measuring for either Malpighian tubules or mouse kidney tubules (Bhave et al., 2017; Howard et al., 2019). Inhibiting Peroxidasin with PHG did not visibly alter Malpighian tubule basement membranes, as assessed by EM (Fig. 7A,B), and their thickness was not changed (Fig. 7C). As predicted from our previous knockdown experiments, pharmacologically inhibiting Peroxidasin for 5 days was sufficient to significantly reduce tissue tensile stiffness by about half as determined from the slope of the stress-strain response at 10-20% strain. These results, combined with previous knockdown results, demonstrate that Peroxidasin is needed in adults to maintain tissue stiffness.

## Conclusions

Analyzing new fly and mouse mutations, we have shown that Peroxidasin mutants in both species are semi-lethal, with variable rates of escaping survivors. We note that mouse viability was assessed at weaning, whereas fly viability was assessed at adulthood, and this may contribute to the apparent reduced viability of fly vs. mouse (13% survival to fly adulthood; 50% survival to mouse weaning). Because even catalytic null mutations show some level of NC1 crosslinking, we conclude that there are other less-specific mechanisms that inefficiently crosslink the NC1 domain. It is likely that these inefficient crosslinking activities ameliorate the loss of crosslinking phenotype and so result in incomplete lethality. Further, the fly Pxn11 mutants appear to be under heavy selection to accumulate modifiers, which may also contribute to the viability of otherwise lethal mutants. Analysis of the developmental lethality of the fly mutants, as well as analysis of their tissue mechanics, demonstrate that Peroxidasin is required for tissue stiffness under and tensile conditions. Finally, the requirement for Peroxidasin in maintaining adult tissue stiffness is surprising because the sulfilimine crosslink that Peroxidasin generates is a covalent bond without any clear mechanism for reversal.

However, these results imply that either the bond must be replaced, or the collagen IV must be replaced over a period of 5 days. The findings of our study reinforce the idea that NC1 crosslinking is essential for animal life.

## Acknowledgements

We are grateful to Hui-Yu Ku for help with the oocyte bursting assay. Funding was provided by F31GM148021 to AMS, R01GM137595 to APM, NSF CAREER 2216394 to NF and XXXX to GB.

## Author Contributions

Conceptualization: KEP, KSL, PSPM, AMS, NF, GB, APM

Methodology: KSL, PSPM, NF

Formal analysis: KEP, KSL, PSPM, DW, NF

Investigation: KEP, KSL, CM, SC, AMS, DW, PSPM

KEP: Fly hypomorphic viability, developmental survival analysis, pupal development, muscle defect

KSL: Pxn mutant generation, null viability, sterility, oocyte bursting

CM: K01 and K02 viability

PSPM: Fly Western

SC: Mouse western, K01 and K02 viability

AMS: EM analysis, dissections for mechanical measurements

DW and NF: mechanical measurements and analysis

Writing: APM, NF, PPM

Visualization: APM, PSPM, NF, KEP

Supervision: APM, GB, NF

Project Administration: APM, GB

Funding Acquisition: APM, GB

## Methods

### *Pxn^11^* deletion

The *Peroxidasin* gene locus was targeted for mutagenesis using two guide RNAs gRNA9: GAAGCAGATCAATTCCCATT gRNA12: TGAACAGTTCCGCAGACTTC pCFD5 plasmids were generated with each gRNA and injected into yw; nos-Cas9 (II-attp40) by BestGene Inc. Injected embryos were crossed to remove Cas9 and were screened by PCR for deletions in the locus, and suspected deletions were sequenced. The Pxn11 deletion removes 727 base pairs, so that the predicted amino acid sequence ends at residue 1040, and asparagine (N) 1041 is gone. After the deletion, the last 2 bp of arginine (R) 1283 are present generating a frame shift which results in eight nonsense mutations, the first of which occurs between leucine (L) 1297 and threonine (T) 1298.

### Analysis of fly lethality through development

25-50 *Pxn^11^/TM3,sChFP* females were crossed with half as many *Pxn^11^/TM3,sChFP* males and placed on a grape juice plate in an egg collection cage at room temperature for one hour before moving them to a new plate supplemented with wet yeast paste for 2 hours at 25°C. After removed parents, embryos were allowed to age overnight at 18°C until roughly stage 13-15. Embryos were sorted by red fluorescence and embryos with visible development were placed on new grape plates supplemented with wet yeast paste, up to 50 embryos per plate. Every subsequent day, progeny were transferred to new grape plates supplemented with wet yeast paste and progeny were counted and developmental stage observed. At pupariation, progeny were transferred from plates to fresh vials with molasses cornmeal food to eclose and were observed daily, including counting adults. Adult phenotype was confirmed to match the appropriate genotype. The percent lost at each stage only included progeny observed at both developmental stages. For assessing pupal development, *Pxn* heterozygotes were crossed and the progeny allowed to develop in the vial. After day 6 when pupae became visible, individual pupae were tracked daily through eclosion and noted for body shape and eye color. Grape plates were made by autoclaving 700 mL dH2O and 30g agar, cooling it slightly, mixing in 300 mL Welch’s grape juice and 20 mL 2.5% Tegosept, and pouring the mixture into 60 mm Petri dishes to cool, stored at 4°C. Wild-type controls were *w^1118^*.

### Analysis of mouse lethality

The *K02* allele was generated from the K01 allele by crossing sequentially to Flp and Cre recombinase lines, thus permanently eliminating exon 9 from the genome. Outcrossing to both the 129S2 or C57/B6 lines was done for ten generations. Genotypes of progeny were determined by PCR of tail clips of pups post-weaning.

### NC1 Western blotting

50 frozen flies of two genotypes, *Pxn^11^* or *w^1118^*, were ground in liquid nitrogen in microfuge tubes then sonicated in 800 µl DOC buffer (10 mM Tris-Cl pH 7.5, 1 M 1% sodium deoxycholate, 1 mM EDTA-Na, pH 8) vortexed and ground again, then incubated on ice with occasional vortexing for 30 min. Tubes were spun 4° for 30 min in a microfuge, and pellets were resuspended in 800 µl collagenase buffer (100 mM NaCl, 50 mM Tris-Cl pH 7.5, 5 mM CaCl2, 25 mM aminocaproic acid, 5 mM Benzamidine) on ice, then vortexed, spun again at 4°, resuspended in 100 µl collagenase buffer. 15 µl collagenase was added from a freshly thawed aliquot, mixed, then incubated overnight at 37°. This collagenase digest was spun 40 min at full speed at 4°, and the supernatant was removed with a gel-loading fine tip pipet to a new tube. Any remaining debris was spun out and supernatant was mixed with 1X sample buffer without DTT, boiled 60 min, analyzed by SDS-PAGE, transferred to nitrocellulose and allowed to air-dry overnight at RT, then re-wet in PBS. Primary antibody as rabbit anti-fly NC1 (1:500) (McCall et al) and LICOR anti-rabbit prepared according to manufacturer’s instructions. About 1.9 fly equivalents are loaded in the 1X lanes.

### Gut imaging

Optical cross sections of the adult Drosophila posterior midgut were obtained as previously described in Howard et al.

### Oocyte bursting

*Pxn^11^* and *w^1118^* females were aged 5-7 days at 25° C on normal food. Ovaries were dissected in cold Schneider’s media then transferred to a glass microscope slide with a 26 x 16 x 0.15 mm spacer (iSpacer from SunJin Lab) containing 75 µl water, then covered with a 24 x 40mm cover slip. Movies were taken with 5x objective on Zeiss Apotome M2 for 10 minutes, with images taken every 3 seconds.

### Electron microscopy

Malpighian tubule samples were processed for TEM and imaged in the Vanderbilt Cell Imaging Shared Resource-Research EM facility, as described in Howard et al., 2019. Basement membrane thickness was measured by applying a grid over the image in FIJI and measuring the thickness of the basement membrane each time the grid intersected. Each data point represents a mean of 10–70 replicate measurements.

### Stiffness measurements of Malpighian tubule basement membrane

Flies were fed 100 µM PHG or water equivalent mixed in 0.2 g Gerber peach baby food in an Eppendorf tube cap. The cap of food was taped into an egg laying cage, and 15-20 aged and mated female flies were added along with a few males. The cages were placed in a humidified chamber at 29 °C. The food was replaced and the chambers were washed daily for 5 days. Malpighian tubules were microdissected and then transferred to cold PBS. The stiffness of native and phloroglucinol (PHG) treated Malpighian tubule basement membrane was characterized using a cantilever-based tensile stiffness measurement system described previously (Sant et al., 2020). We showed in prior studies that the cellular contribution to the overall tubule stiffness is minimal, so the measured stiffness represents the stiffness of basement membrane (Bhave et al., 2017; Howard et al., 2019). Cantilever stiffness sensors were fabricated from glass capillary tubes using a pipette puller and cantilever spring constants were calibrated in a manner similar to Shimamoto and Kapoor (Shimamoto and Kapoor, 2012). Malpighian tubules were attached to the cantilever and a rigid holding pipette, shown in Fig. 7. The movement of the holding pipette was controlled by a micromanipulator at a speed of 10 μm/sec to stretch the tubule and imaged in increments of 40 μm displacements. Images were captured with a digital camera attached to an inverted microscope (VWR) at 4× magnification. Displacement of the cantilever and holding pipette were analyzed in WINanalyze software. The stress was calculated as the applied force divided by the cross-sectional area of the basement membrane. The average width of the basement membrane used to calculate the cross-sectional area as measured from TEM images and tubule diameter was measured by light microscopy (Bhave et al., 2017; Howard et al., 2019). To model the stress-strain response, Humphrey model was employed to fit the experimental data. The stretch ratio was taken as the ratio of the deformed tubule length to the initial length (Sant et al., 2021; Sant et al., 2020) and the strain = stretch ratio-1. The equations for the hyperelastic model were used in the functional form described by Martins et al (Martins et al., 2006). The Humphrey model was fit to the experimental data using lsqcurvefit function and the elastic moduli were calculated from the slope of the stress strain response for strain from 10-20%. Representative curve fits to the experimental data and the model fit parameters are provided in the supplementary data (Figure S3).

---

**Figure S1:**
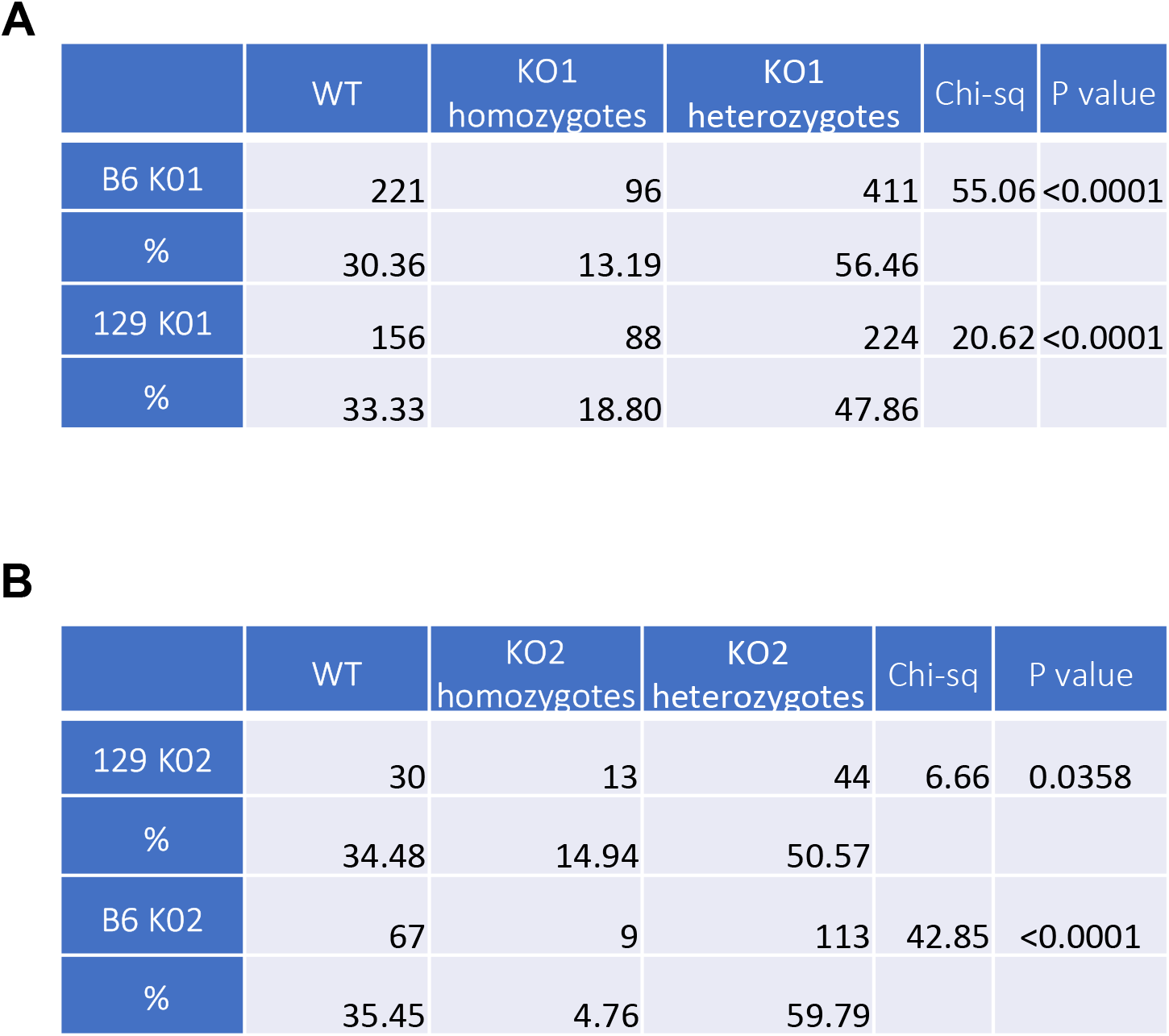
Chi-square and p values for K01 and K02 homozygote viability.

**Figure S2:**
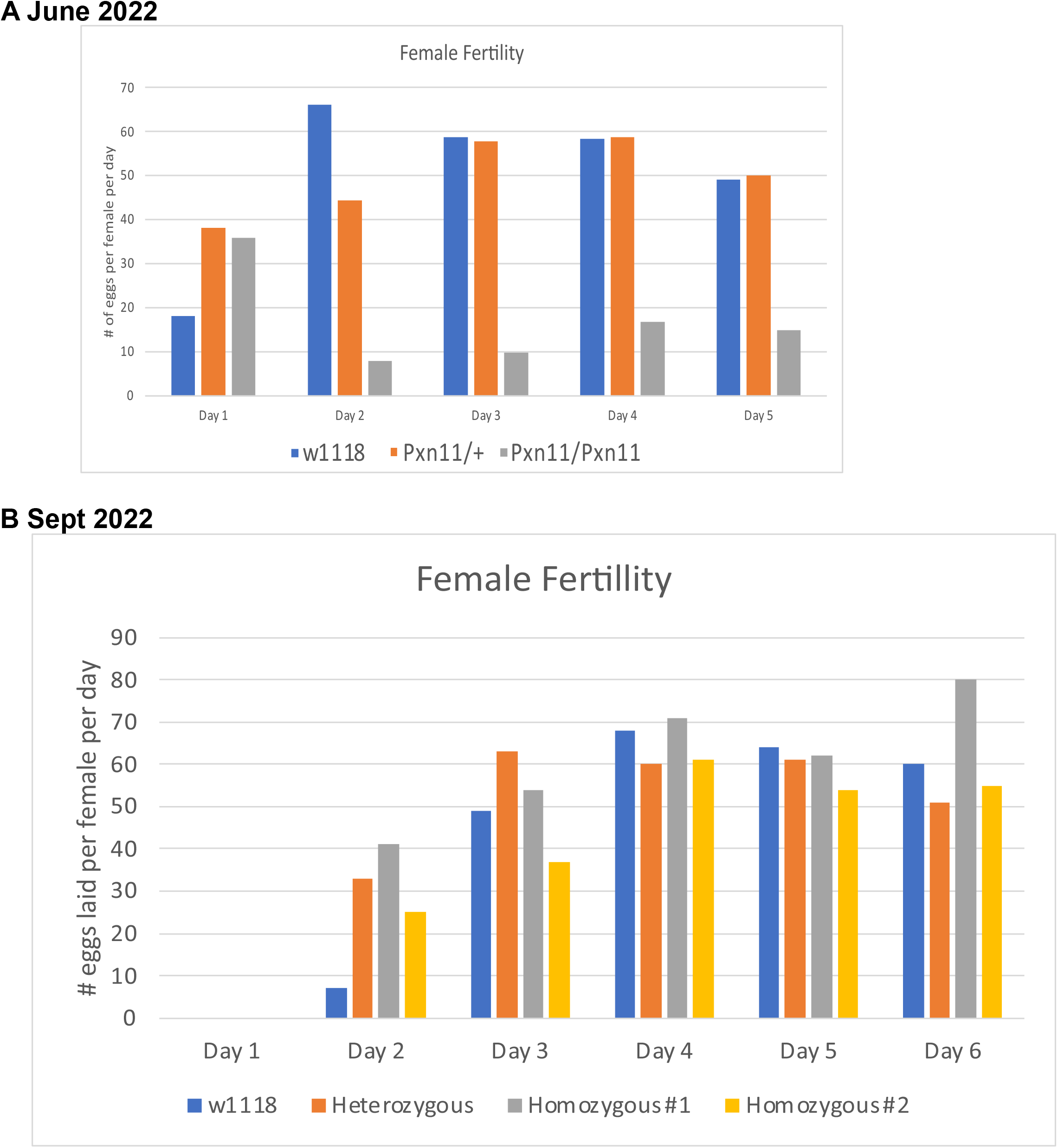
Fertility of *Pxn¹¹* mutant females increased over months, suggesting the presence of modifiers. **A.** In June 2022, *Pxn¹¹* mutants displayed severely reduced egg-laying capacity compared to homozygous wild-type and *Pxn¹¹* heterozygous controls. Fertility was measured as number of eggs laid per day divided by the number of females. **B.** In Sept. 2022, two populations of *Pxn¹¹* mutants (#1 and #2) displayed normal egg-laying, similar to wild-type and *Pxn¹¹* heterozygous controls.

**Figure S3:**
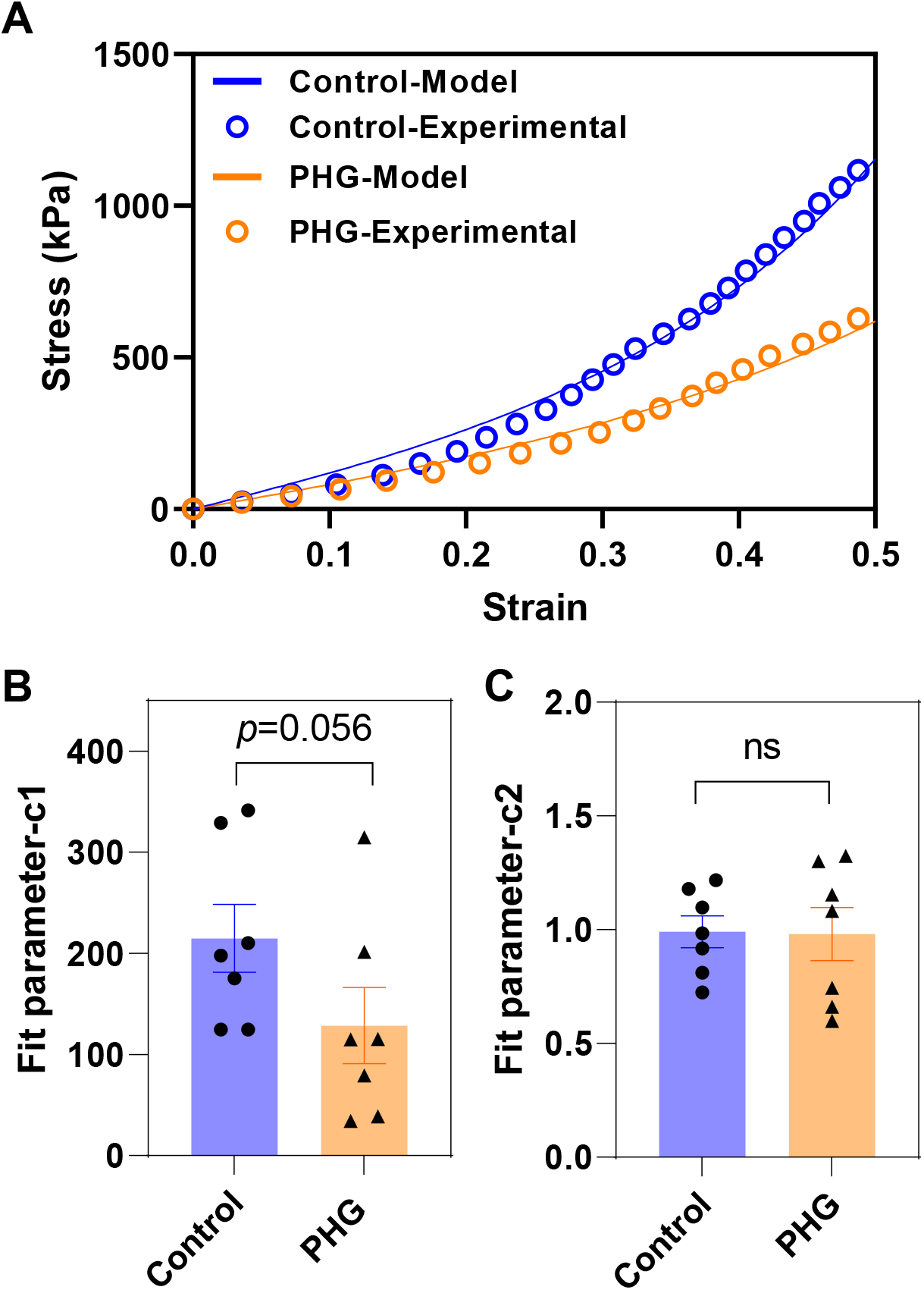
Experimental stress-strain data fit to a Humphries model of hyperelastic materials. **(A)** Representative strain-stress experimental data and model fit curves of control and PHG treated Malpighian tubules. **(B, C)** Comparison of the model fit parameters, c1 and c2, between control and PHG treated Malpighian tubules. Data were analyzed by unpaired t-test in GraphPad, n=7 for all conditions.

